# *Bordetella* oligosaccharide (BOS) is associated with lipopolysaccharide of *Bordetella petrii* - the ancestor-related species of the pathogenic *Bordetella*

**DOI:** 10.64898/2026.04.29.721566

**Authors:** Sabina Koj, Karolina Ucieklak, Olga Rojewska, Tomasz Niedziela

## Abstract

*Bordetella* produce a wide array of virulence factors. These factors are involved in bacterial colonization and evasion of immune defenses. Our recent studies revealed that the bacteria produce an exoglycan, *Bordetella* oligosaccharide (BOS). *B. petrii* is the evolutionary early divergent species of the genus *Bordetella*. This study has focused on the investigation of two *B. petrii* type strains: clinical and environmental. We employed nuclear magnetic resonance (NMR) analyses to elucidate the structural differences between their lipopolysaccharides. Our findings revealed that the LPS of clinical *B. petrii* strain comprises a hexasaccharide unit, that was structurally identical to the BOS. This form of LPS is only a minor population in the bacterial outer membrane of the environmental strain. In addition to the cell-bound BOS, its secreted glycoform was also found in growth media of *B. petrii*. Anti-BOS neoglycoconjugate antibodies cross-reacted with *B. petrii* LPS. This suggest that the newly identified BOS associated with *B. petrii* PS would be a potential vaccine element against *Bordetella*.

## Introduction

The *Bordetella* genus consists of sixteen species, which include pathogens responsible for respiratory infections. *B. pertussis* and *B. parapertussis* cause whooping cough in humans while *B. bronchiseptica* mostly infects animals ^1^. The other species of *Bordetella* comprise *B. holmesii* ^2,3^, *B. trematum* ^4,5^, *B. ansorpii* ^6^, *B. avium* ^7^, *B. hinzii* ^7^, *B. petrii* ^*8*^, *B. bronchialis, B. flabilis*, and *B. sputigena* ^9^. Recently, *B. hinzii*-like isolates from laboratory-raised mice have been delineated and classified to *B. pseudohinzii* ^10^. Whereas, *B. muralis, B. tumulicola*, and *B. tumbae* are newly recognised environmental species isolated from 1300-year-old mural paintings in Japan ^11^. The latter ones are closely related at the molecular and phylogenetic level to *B. petrii*, which previously had been described as the only species with an environmental origin found among the otherwise host-restricted and pathogenic members of the genus *Bordetella* ^12^.

The genomes of some *Bordetella* have been sequenced, providing insight into the evolutionary history of the genus and closely related groups, that are *Achromobacter* and *Alcaligenes* within the *β-Proteobacteria* ^12–14^. It has been suggested that *B. pertussis* and *B. parapertussis* have evolved independently from a *B. bronchiseptica*-like ancestor ^15^, whereas the genome sequence of *B. petrii* revealed divergence from the typical *Bordetella* pathogens. The environmental isolate, *B. petrii* strain DSMZ 12804 has a mosaic genome of 5,287,950 bp with numerous mobile genetic elements, including genomic islands highly related to a *clc* element of *Pseudomonas knackmussii* B13, which encodes genes involved in the degradation of aromatics. The *B. petrii* also express virulence factors, such as a filamentous hemagglutinin, and the master virulence regulator BvgAS ^12^, but it lacks toxins of the pathogenic *Bordetella*. The evolution of the human-adapted *Bordetella* species has been accompanied by massive genome reduction, lack of horizontal acquisition of genetic material and significant differences in virulence gene expression among species ^13,16^. Furthermore, the adaptation to the specific organism may have caused a reduction of the properties required for survival in the environment and involved in metabolic functions including biosynthesis of lipopolysaccharide, amino acid metabolism and degradation of aromatic compounds ^17^. Therefore, the genomic analysis suggests that phylogenetically *B. petrii* connects the host-restricted obligate pathogenic *Bordetella* and free-living environmental bacteria of the genera *Achromobacter* and *Alcaligenes* ^12^.

*B. petrii* strains were repeatedly found in different environmental niches, such as polluted soil, river sediments, waste water, marine sponges, rhizosphere and a grass root ^18,19^. The presence of heavy metal resistance systems and remarkable metabolic versatility enables *B. petrii* to thrive in the different ecosystems ^20^. Furthermore, *B. petrii* also exhibits properties of host-restricted pathogen, and is well-adapted to the human host. The clinical isolates of *B. petrii* have been associated with bone degenerative disease, mandibular osteomyelitis ^8^, mastoiditis ^21^, chronic pulmonary disease ^22^ and cystic fibrosis ^23^.

*Bordetella* are Gram-negative bacteria and lipopolysaccharide (LPS) is the main component of their bacterial envelope. The structures of the LPS vary from complete LPS through semi-rough LPS, to O-antigen deficient lipooligosaccharide (LOS). O-specific polysaccharide (O-PS) components of *B. bronchiseptica* and *B. parapertussis* LPS are composed of linear polymers of 1,4-linked 2,3-diacetamido-2,3-dideoxy-α-L-galactopyranosyluronic acid residues ^24^. Some of these sugars are found in the uronamide form, while the terminal residues of the O-PS contain unusual modifications ^25^. Subunits with characteristic diaminouronic acid derivatives were identified also in O-chains of *B. avium* ^26^, *B. trematum* ^27^ and *B. hinzii* ^28^. In contrast, *B. pertussis* produces LOS, which consists of a lipid A moiety linked to a core dodecasaccharide that contains a distal trisaccharide built of N-acetylglucosamine, 2,3-diacetamido-2,3-dideoxy-mannuronic acid, and 2-acetamido-4-N-methyl-2,4-dideoxy-fucose. Some strains of *B. pertussis* possess a core nonasaccharide devoid of the distal trisaccharide. Such core represents the conserved element of *Bordetella* LPS ^29^ and is shared in most of *Bordetella* ^30^.

So far, lipid A was the only structurally characterized part of *B. petrii* LPS. The *B. petrii* lipid A differs in the length and degree of acylation depending on the environmental or human origin of the isolates. One of the unusual features of the lipid A is the absence of symmetry at the C-3 and C-3′ positions and the presence of a short-chain 10:0(3-OH) at the C-3 position, that is more characteristic for the human pathogens ^19^. LPS is the major immunogenic component of *B. petrii*, according to literature ^23^. The isolates with a deletion in glycosyltransferase within the putative O-antigen biosynthesis locus, leading to O-antigen loss are more susceptible to a complement-mediated killing. Except lipopolysaccharides, bacterial pathogens produce several types of polysaccharides vital for colonization and pathogenesis that provide protection against the host defense mechanisms. Among them, capsular polysaccharides (CPS) are bound to the outer membrane of the bacteria, whereas the extracellular polysaccharides (EPS) are secreted out of the bacterial cell. The presence of extracellular glycans has been predicted from genomic analyses of *Bordetella*. According to the literature data, the major polysaccharide produced by *B. pertussis* is *Bps* (*Bordetella* polysaccharide). With a PNAG-like structure, the *Bps* is antigenically and biochemically similar to the poly-β-1,6-*N*-acetylglucosamine ^31^, intercellular adhesin originally isolated from the biofilm of *Staphylococcus aureus* that plays a significant role in colonisation, biofilm formation, and enhances resistance to antimicrobials and phagocytosis ^32–35^. Another exopolysaccharide reported for *B. pertussis* is a linear polymer composed of N-acetylated galactosaminuronic acid residues that is similar to Vi antigen, found previously in *Salmonella enterica* serovar Typhi ^36^. Vi antigen reduces serum complement killing and promotes intracellular replication. The genetic locus for Vi antigen is present in *Bordetella* although the glycan product has been identified only serologically. In our parallel studies, we have identified a new glycan in *Bordetella* bacterial cultures. This extracellular molecule universal among *Bordetella* structurally is a hexasaccharide, therefore it was named *Bordetella* oligosaccharide (BOS) ^37^. However, biological significance of this new type of glycan remains unknown.

Most *Bordetella* have adopted to human host in the evolutionary process of the genera. The selective pressure from the host”s immune defense could have driven antigenic variability of *Bordetella* O-PS, that allowed the bacteria to evade a cross-reaction with other *Bordetella* and to escape from the host immunity. With a high genetic diversity and metabolic flexibility, *B. petrii* exhibits a broad adaptability. *B. petrii*, the opportunistic human pathogen, has environmental origin and represents a lineage that diverged early, therefore we presume that this bacterium exhibits the LPS with structure distinct from other *Bordetella*, and with additional structural features. Here, the studies include structural comparison of the primary clinical strain of *B. petrii* NCTC 13363 and environmental isolate of *B. petrii* DSM 12804, using NMR spectroscopy and mass spectrometry. The virulence factors may differ depending on environmental or human origin of the strains. Investigation of the early diverged lineage of *B. petrii* might reveal the pathogenic molecular patterns shared among *Bordetella*, important as vaccine targets.

## Results

### *B. petrii* LPS and oligosaccharide fractions

*B. petrii* strains NCTC 13363 and DSM 12804 were grown in the Stainer-Scholte medium and subsequently LPS were extracted from bacterial cells by the hot phenol/water extraction method ^38^, and purified by ultracentrifugation. The yields of the LPS were estimated in relation to dry bacterial mass - 0.6% (*B. petrii* 13363) and 0.32% (*B. petrii* 12804). LPS isolated from water phase was further investigated. The SDS-PAGE analysis of the *B. petrii* LPS was performed and the profiles were compared to this of LOS of *B. pertussis* 186 (Fig. 1 A). *B. pertussis* 186 LOS comprises a core oligosaccharide substituted with a distal trisaccharide, which appears as the higher molecular weight “A band” on SDS-PAGE. The “B band” is minor fast-migrating band representing a smaller, incomplete LOS lacking this terminal trisaccharide. The analysis of *B. petrii* 12804 LPS showed “B band”, whereas molecular size of *B. petrii* 13363 LPS is higher than the size of *B. pertussis* 186 LOS. The molecular mass differences in the SDS-PAGE profiles suggest the presence of additional element in the *B. petrii* 13363 LPS.

**Fig. 1.**
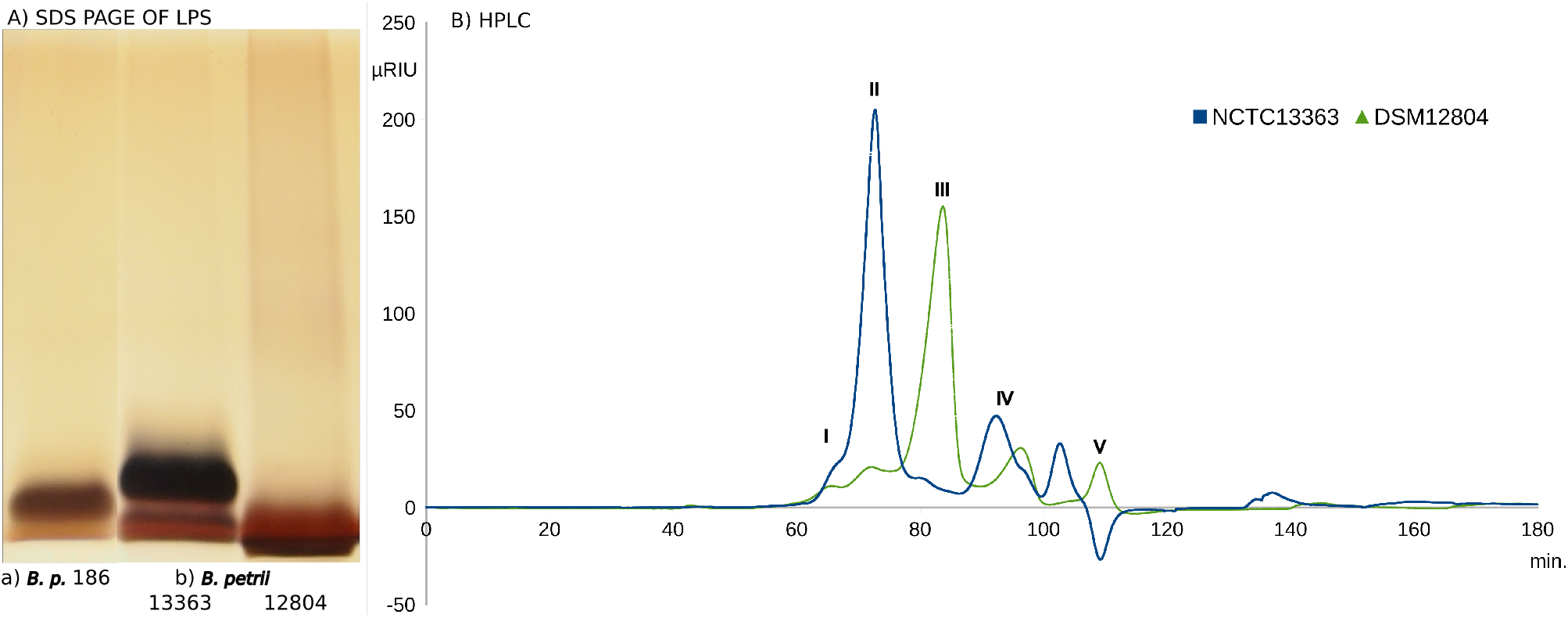
Analysis of *Bordetella* LPS and isolated glycans. A) SDS-PAGE profiles of LPS from a) *B. pertussis* 186, b) *B. petrii* NCTC 13363 and *B. petrii* DSM 12804. B) Gel-filtration chromatography profiles of the heterooligo- and polysaccharides isolated from *B. petrii* 13363 and *B. petrii* 12804 LPS, separated on the Superdex 30 pg with RI detection. The fractions I-V are indicated.

The isolated *B. petrii* LPS of strains 13363 (36.8 mg) and 12804 (36.2 mg) were hydrolyzed with 1.5% acetic acid. Lipids A were separated and the soluble products were further fractionated by size-exclusion chromatography (Superdex 30 pg). In the profiles, the five fractions were distinguished according to the retention times: I 58-66 min, II 67-78 min, III 79-90 min, IV 91-105 min and V 106-115 min (Fig. 1 B). The main peaks for the *B. petrii* strains differed. The main fraction isolated from *B. petrii* 13363 hydrolysate was the fraction II (6 mg), while the fraction III (7.2 mg) represents the major population of *B. petrii* 12804 OS. The high molecular weight fraction II (1.4 mg) of *B. petrii* 12804 was a minor population of the heteroligosaccharides.

### *B. petrii* core oligosaccharides

The main fractions, III of *B. petrii* 12804 and II of *B. petrii* 13363 were analyzed using NMR spectroscopy. The anomeric regions of the ^1^H NMR and ^1^H, ^13^C HSQC spectra of both fractions indicated 7 signals with similar chemical shifts (Fig. 2). Comparative analysis with the published data indicated that the ^1^H and ^13^C chemical shift values correspond to the residues of the inner core oligosaccharides of other *Bordetella* ^29^. According to the data, the anomeric signals were assigned as residues A-H (Fig. 2 A), without the E residue. The characteristic signals indicative of a terminal trisaccharide, e.i. signal corresponding to the N-methyl of Fuc2NAc4NMe, were absent. Further analysis using ^1^H, ^1^H and ^1^H, ^13^C correlation experiments, including ^1^H, ^1^H COSY; ^1^H, ^1^H TOCSY; ^1^H, ^1^H NOESY, ^1^H, ^13^C HMBC and ^1^H, ^13^C HSQC-TOCSY confirmed that *B. petrii* 12804 core oligosaccharide is devoid of terminal α-Glc*p*N, while 2-substituted L-α-D-Hep*p* (residue B) is decorated with phosphate group at the C-4 position (Supplementary Fig. 1, Supplementary Table 1). With the 2-substituted L-α-D-Hep*p*4P, the fraction III of *B. petrii* 12804 constitutes the inner core part, comprising subsequently the residues α-Gal*p*NA (D), α-Glc*p*A (G), L-α-D-Hep*p* (C), 4-substituted α-Glc*p*N (A), 4,6-disubstituted β-Glc*p* (H), 3,4-disubstituted L-α-D-Hep*p* (F) and Kdo (I and I”). The residue F was present in two variants, F and F”. By the initial comparative analysis, ^1^H NMR spectrum of fraction II suggests that the residues of the core oligosaccharide are present also in the *B. petrii* 13363 PS (Fig. 2 B). Moreover, the spectrum of the *B. petrii* 13363 fraction II except the signals corresponding to the core, contains six additional anomeric signals and intensive signals from acetyl groups. The additional hexasaccharide element was further investigated to determine its structural features and identify its linkage to the core oligosaccharide.

**Fig. 2.**
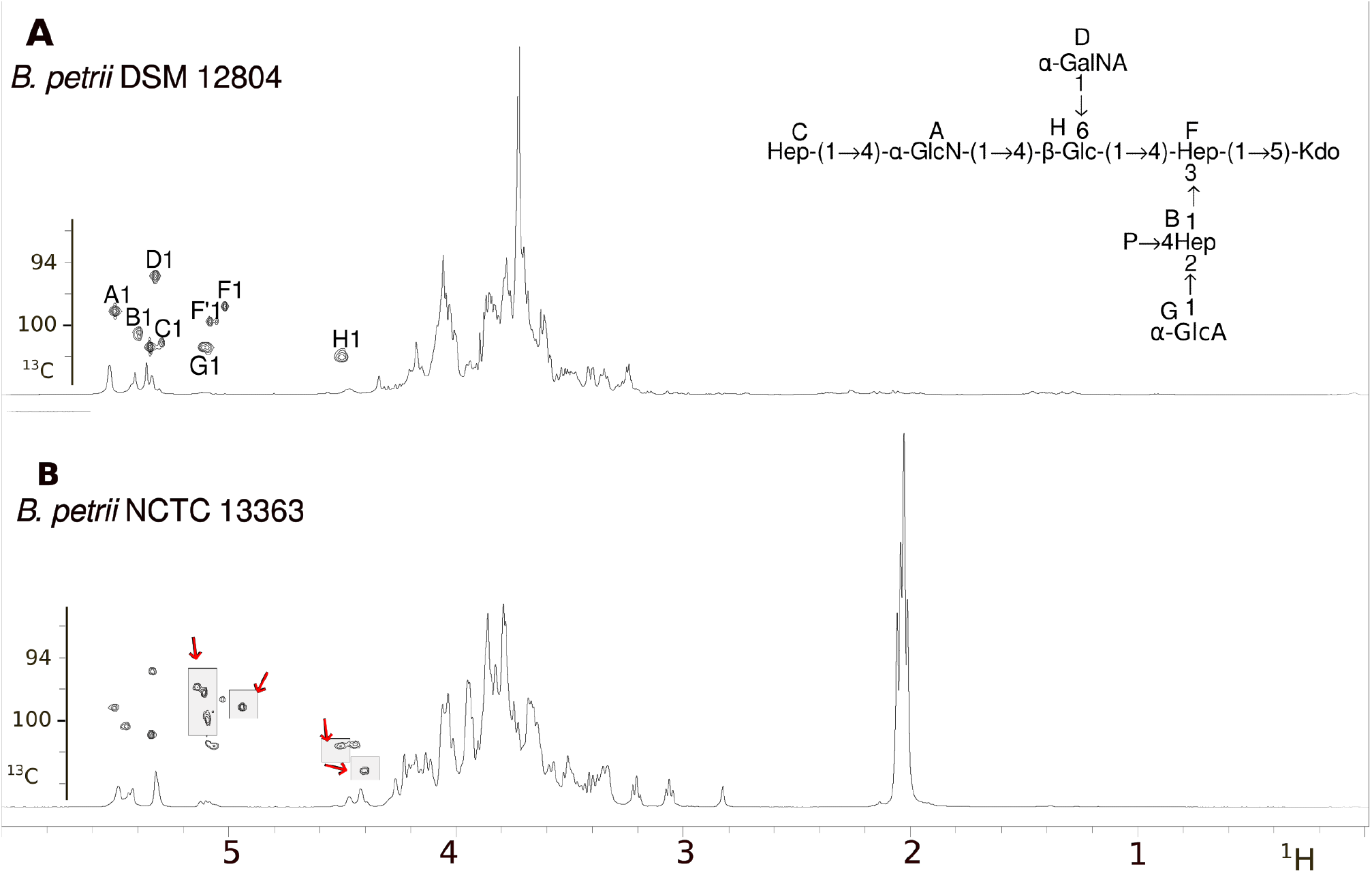
NMR analysis of *B. petrii* glycans. ^1^H NMR spectra and anomeric regions of ^1^H, ^13^C HSQC spectra of: A) *B. petrii* 12804 fraction III and B) *B. petrii* 13363 fraction II. The shared signals are assigned as residues A to H, whereas E residues is absent in the spectrum of fraction III. The additional signals in the spectra of *B. petrii* 13363 fraction II are indicated in the gray boxes.

### *B. petrii* hexasaccharide unit and its linkage to the LPS core

The ^1^H, ^13^C HSQC-DEPT, HSQC-TOCSY and HMBC spectra of the *B. petrii* NCTC 13363 fraction II contains 14 anomeric signals (Fig. 3 A and Supplementary Fig. 2, Table 1). Some of the signals correspond to the oligosaccharide that resemble the inner OS of *Bordetella*, as was indicated in Fig.

**Fig. 3.**
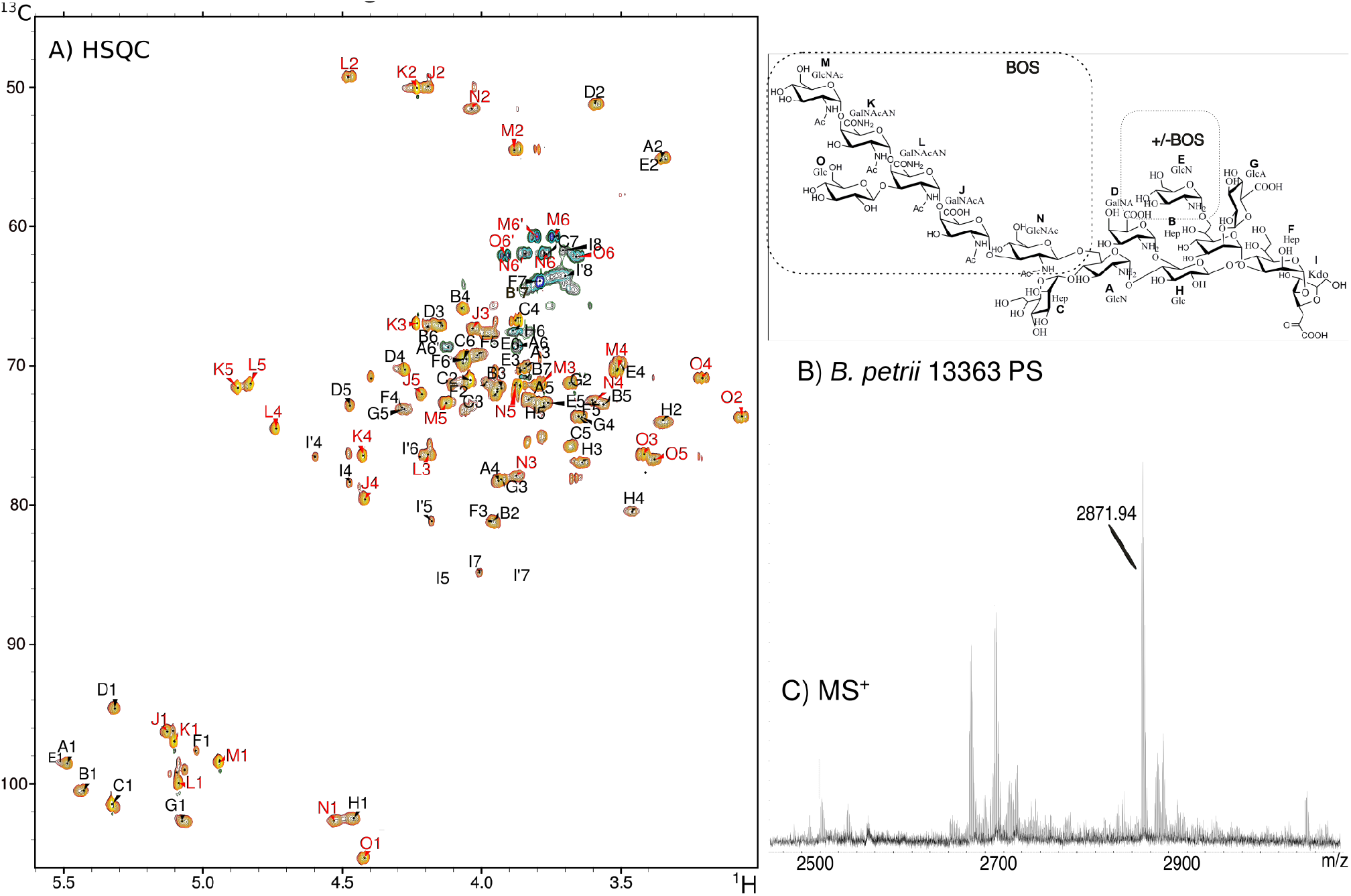
Structural characteristic of *B. petrii* NCTC 13363 PS. A) ^1^H, ^13^C HSQC spectrum of fraction II. Signals indicating the BOS unit of the *B. petrii* PS are shown in red. The capital letters refer to carbohydrate residues as shown in B) structure of *B. petrii* 13363 PS - core OS with one hexasaccharide unit linked to the residue A. The PS shows heterogeneity - the residue E can be substituted with the second unit of the BOS. C) MALDI-TOF MS of the *B. petrii* 13363 PS recorded in the positive mode with DHB as a matrix. The main ion corresponds to the PS structure.

**Table 1.**
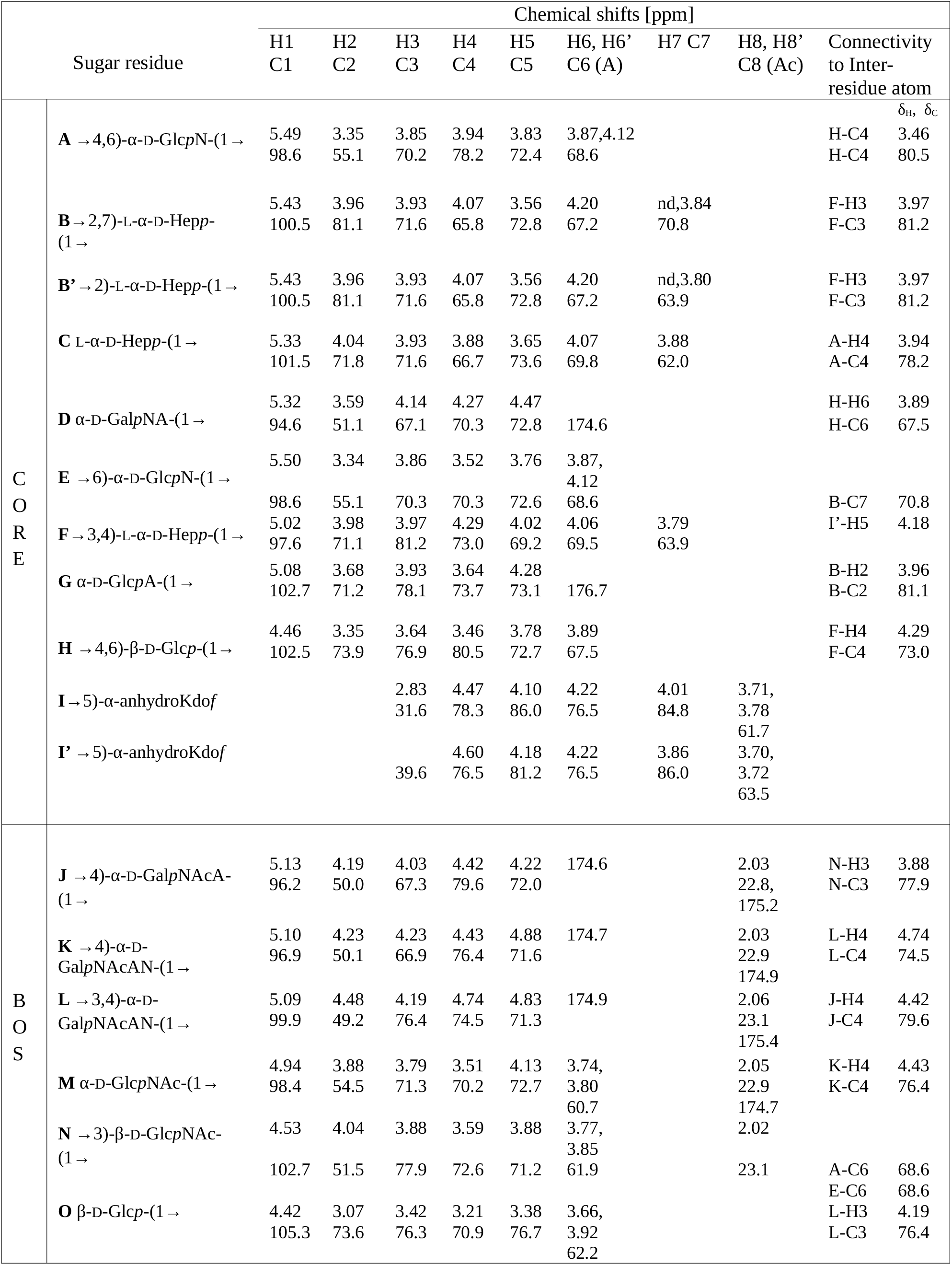
^1^H and ^13^C chemical shifts of fraction II from *B. petrii* 13363 PS. nd – not determined.

2. The residues **A**-**I”** constitute the core nonasaccharide of the *B. petrii* LPS that comprises the L-α-D-Hep*p* (C), α-Gal*p*NA (D), α-Glc*p*A (G), 4,6-disubstituted β-Glc*p* (H), 4,6-disubstituted α-Glc*p*N (A), 6-substituted α-Glc*p*N (E), 2,7-disubstituted L*-α-*D*-*Hep*p* (B), 3,4-disubstituted L-α-D-Hep*p* (F) and 5-substituted anhydroKdo residues (I and its variant I”). Formation of the 4,7-furanoid derivatives of Kdo was a result of a mild acid hydrolysis and the β-elimination of the phosphate from the C-4 position of the 5-substituted 2-keto-3-deoxy-D-*manno*-octulosonic acid in the native PS ^39–41^. The 2-substituted B” variant was also identified, indicating microheterogeneity from a presence of the PS glycoform devoid of E residue.

The remaining spin systems of the **J**-**O** residues correspond to a hexasaccharide that includes the α-GlcNAc (M), β-Glc (O), 4-substituted α-GalNAcAN (K), 3,4-disubstituted α-GalNAcAN (L), 4-substituted α-GalNAcA (J) and 3-substituted β-GlcNAc (N) residues, with the following sequence MK(O)LJN. The N residue is partially N-acetylated. This hexasaccharide component of *B. petrii* 13363 LPS, is structurally identical to the recently identified exo-oligosaccharide of *Bordetella* (BOS), which structure was described previously ^37^.

This BOS unit substitutes the core oligosaccharide of the *B. petrii* 13363 LPS. In the BOS_LPS_, the 3-substituted residue N is the linkage point of the BOS to the core OS - through the A residue at the position 6. In result, the structure of the *B. petrii* 13363 PS composed of the core oligosaccharide substituted by the BOS was determined with the following sequence: α-GlcNAc-(1→4)-α-GalNAcAN-(1→4)-[β-Glc-(1→3)]-α-GalNAcAN-(1→4)-α-GalNAcA-(1→3)-β-GlcNAc-(1→ →6)-[Hep-(1→4)]-GlcN-(1→4)-[GalNA-(1→6)]-Glc-(1→4)-{GlcA-(1→2)}-Hep-(1→3)]-Hep-(1→5)-Kdo

The chromatographic profile of the heteropoly- and oligosaccharide mixture of the *B. petrii* 13363 PS indicates a dominant population of the fraction II - a core substituted with a single hexasaccharide unit, with a minor heterogeneity resulting from the presence of one fraction with higher molecular weight (fraction I) and three fractions with the lower molecular sizes (fraction III, IV and V). In a spectrum of the fraction I, the signals corresponding to the hexasaccharide, and relatively weaker signals from the core suggest the presence of the oligosaccharide with two hexasaccharide units, attached to both glucosamine residues, A and E at the position 6 (Fig. 3 B, Supplementary Fig. 3, Supplementary Table 2). The fractions III and IV contained the unsubstituted core OS (Supplementary Fig. 4, Supplementary Table 2). Fraction V contained α-glucosamine phosphorylated at C-1 (P→α-GlcN). The spectra of the unfractionated *B. petrii* 13363 PS contains the glucosamine-1-phosphate and different variants of glucosamines that might confirm the role of the GlcN residues as the linking points of the hexasaccharide to the core (Supplementary Fig. 5).

The chromatographic profile of *B. petrii* 12804 heterooligosaccharides indicates a dominant population of the fraction III comprising the core oligosaccharide. Whereas, the NMR spectra of the minor fraction II contained the signals of the core OS and weak signals from the hexasaccharide, suggesting that the core substituted with the BOS unit is present also in *B. petrii* 12804 OS (Supplementary Fig. 6), but in relatively low amount (16% of the total population).

To verify the structures of *B. petrii* poly- and oligosaccharides, the fractions have been investigated using the MALDI-TOF MS. The mass spectrum of the *B. petrii* 12804 fraction II contained a main ion at *m/z* 1578.80 [M-H_2_O-H, 2Na]^-^, that confirms the structure of the phosphorylated octasaccharide. The MS profile of *B. petrii* 13363 fraction II contained a main ion at *m/z* 2871.94 [M-H_2_O+H, 2Na]^+^ (Fig. 3 C), corresponding to the pentadecasaccharide with anhydroKdo at the reducing end. In the spectrum of the fraction III of *B. petrii* 13363, the main ion at *m/z* 1467.55 [M-H]^-^ corresponded to the core OS devoid of one HexN residue. The mass differences confirmed that the *B. petrii* 13363 PS fraction II is mainly built of the core nonasaccharide and the single hexasaccharide unit.

### *Bordetella* oligosaccharide from *B. petrii* bacterial culture

The presence of the *Bordetella* oligosaccharide (BOS) was revealed in growth media of all investigated *Bordetella, B. pertussis, B. parapertussis, B. holmesii*, and *B. bronchispetica* ^37^. The amounts of the exo-hexasaccharides differed among *Bordetella*, while the average yield was estimated ∼ 2.75 mg per 1 g of the dry bacterial mass.

To verify the presence of the BOS in the growth media of *B. petrii*, the supernatant obtained from the bacterial culture (5 L) was freeze-dried and subsequently the product was fractionated using size-exclusion chromatography (2 g). All fractions were checked by NMR spectroscopy. The fractions were collected according the retention times: fr. I (40-50 min), fr. II (51-69 min), and fr. III (70-82 min). The subsequent fractions contained high amount of salts from culture media. The fraction III was the main fraction, in the profile of the *B. petrii* 13363 (1.17 mg) and of *B. petrii* 12804 (0.77 mg). The chemical shifts of the signals in the spectra of the fraction III were identical to these of the K, L, M, O residues of the hexasaccharide part of *B. petrii* LPS, whereas the values for J and N residues varied - the N residue was present as a reducing end of this glycan (Supplementary Fig. 7, Supplementary Table 2). The new signals were assigned as the N” and N”“ as the 3-linked β-GlcNAc and 3-linked α-GlcNAc, respectively. The modified spin system of J residue was assigned as J” with anomeric chemical shifts δ_H/C_ 5.34/99.5. The presence of the 3-substituted Glc*p*NAc as the reducing end residues confirms that the hexasaccharide is unbound molecule in the *B. petrii* medium. No anchor element was identified in the culture media.

Except the exo-hexasaccharide, the NMR analysis indicated that the growth medium of *B. petrii* contained polymers of glucoses, →4)-α-D-Glc*p*-(1→ and →6)-α-D-Glc*p*-(1→, assigned as the P and R residues, respectively. The glucans were also identified in the culture media of other *Bordetella*, in our previous studies ^37^. The polymers were present in the high-molecular fractions, I and II, whereas the lower-molecular glucan chains were identified also in the fraction III. The distribution of the glucans differed in the fraction I (0.21 mg and 1.85 mg), and fraction II (0.43 mg and 0.15 mg) of the *B. petrii* 13363 and *B. petrii* 12804, respectively. Additionally, the intensive signals corresponding to the uridine diphosphate (UDP) were detected and identified in the fraction III. ^1^H, ^31^P HMBC indicated the correlation of the pyrophosphate to the C1 of NAc-glucosamine and the ribose-C5 of the UDP.

Regarding the relatively high quantities of the exo-hexasaccharide produced by the *Bordetella*, we hypothesized that the role of this glycan is biologically important, therefore the immunogenicity studies were further conducted.

The conjugate of the BOS with a tetaus toxoid as a carrier was prepared and used to immunization of rabbits, in our recent studies ^37^. The obtained anti-hexasaccharide antibodies were cross-reactive in all investigated *Bordetella*. In the present studies, we used the specific anti-hexasaccharide antibodies to reaction with the *B. petrii* NCTC 13363 LPS and *B. petrii* DSM 12804 LPS, in ELISA test. The reactivities of the both *B. petrii* LPS with the anti-hexasaccharide antibodies were similar (Fig. 4 A). Though the fraction containing the hexasaccharide unit represents a minor population of *B. petrii* 12804 LPS, its relatively high reactivity with the anti-hexasaccharide antibodies indicates importance of this epitope within LPS (Fig. 4 B) and specificity of the anti-BOS antibodies. The analysis confirms antigenic identity of the hexasaccharide to the component of the *B. petrii* LPS, as well as to the oligosaccharide secreted to the media during the bacterial culture. Summarizing, the above results allowed us to the conclusions:

**Fig. 4.**
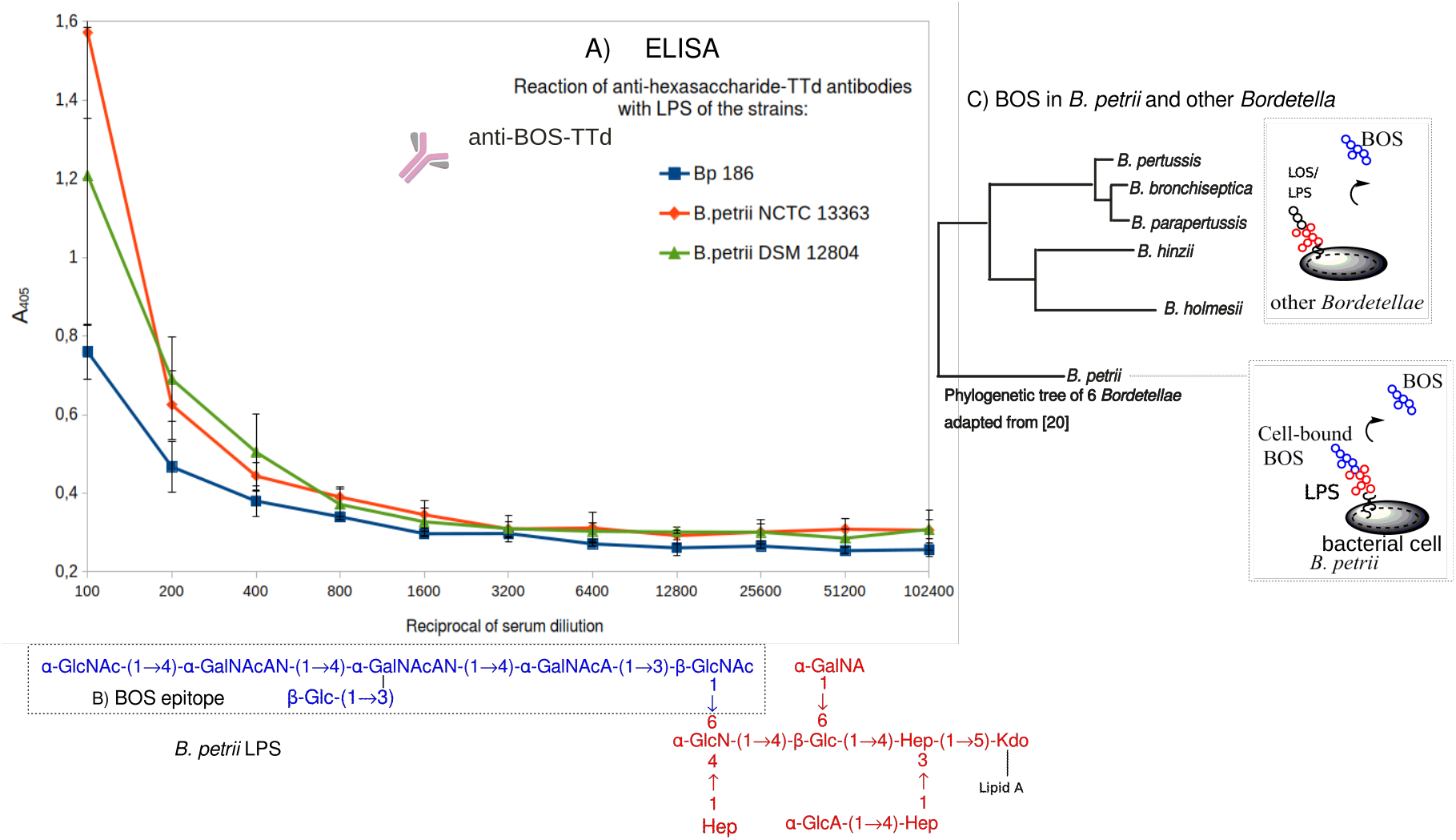
The reactivities of *B. petrii* LPS with the anti-BOS antibodies. A) Reactivities of the polyclonal antibodies raised against the hexasaccharide-TTd conjugate with LPS of *B. petrii* 12804 (green) and *B. petrii* 13363 (red) evaluated by ELISA with the LPS as solid-phase antigen. *B. pertussis* 186 LOS (blue) was used as a control. The data are representative of 3 experiments. B) Epitope of the anti-BOS antibodies within the LPS of *B. petrii*. C) Glycoforms of BOS – cell-bound BOS_LPS_ as well as the released BOS in *B. petrii*, and solely released BOS in other *Bordetella*.

1. The presence of both structurally identical hexasaccharides in *B. petrii*, the LPS-bound glycoform BOS_LPS_ and the free BOS suggests that the BOS originates from LPS.
2. Evolutionary divergence of *B. petrii* from typical pathogenic *Bordetella –* the potential role of the genus progenitor suggests that the BOS could be a component of the LPS in the early evolution of *Bordetella*.
3. Free BOS glycoform present in other investigated *Bordetella* indicates its role in adaptation to host as the pathogenic species (Fig. 4 C).

## Disscusion

Lipopolysaccharide is one of the immunostimulatory components of bacteria and belongs to the pathogen-associated molecular patterns (PAMPs). The structural scheme of LPS is sustained across Gram-negative species, indicating long-term evolutionary importance of LPS. The conserved nature of the bacterial antigens allows the host immune system to recognize and respond to a wide range of bacteria. Simultaneously, evolution pressure from the host immune defense might induce temporary or permanent adaptive changes in bacteria. To adapt to dynamic environments and variety of hosts, evade host immune response and resist killing, pathogens can undergo genetic changes as beneficial mutations. Bacteria have evolved many enzymes that modify outer membranes, by altering properties, charge, permeability, and binding to immune cells. The adaptive evolution might affect the LPS structure, and consequently it may acquire subtle modifications, such as changing acyl chain length of lipid A, adding amine groups to lipid A, or altering the O-antigen ^42^. Some modifications are non-stoichiometric, leading to a microheterogeneous population of LPS molecules that increase variety of the antigen. All the modifications can modulate host immune response, for instance by adding positive charges the overall charge density of LPS molecules is neutralized, impairing a recognition by immune receptors ^42^.

Among all *Bordetella*, the innermost core oligosaccharide domain of LPS is conserved. The core hexasaccharide comprises the α-Glc*p*N, α-Glc*p*A, 4,6-disubstituted β-Glc*p*, 2,7-disubstituted L-α-D-Hep*p*, 3,4-disubstituted L-α-D-Hep*p*, and the Kdo residues and is the least variable OS region of *Bordetella* ^29^. The strains with the rough-type LPS are relatively susceptible to bacterial killing by host immune cells. Whereas, the terminal trisaccharide addition, α-D-Glc*p*NAc-(1→4)-β-D-Man*p*2NAc3NAcA-(1→3)-β-L-Fuc*p*2NAc4NMe-(1→ to the OS core appears important for resistance to pulmonary surfactant protein A ^26,43^. Alternative strategies have evolved in this pathogen to inhibit complement activation, such as an autotransporter protein Vag8, that binds to the C1 esterase inhibitor ^44,45^. Other *Bordetella*, such as *B. parapertussis* typically express protective O-chains, that mediate resistance to the membrane attack complex of the complement pathways. Moreover, we previously described the presence of the additional exo-hexasaccharide revealed in *Bordetella* ^37^. The BOS was found as unbound and structurally simple exo-oligosaccharide in the bacterial external environment that in contrast to the high-molecular weight exopolysaccharides forming complex biofilm matrix in bacteria seemed incomplete. Therefore, the presence of anchor linking the exo-hexasaccharide or multicomponent complex associated with this glycan have been concluded. However, all the attempts to identify such anchor failed.

In the report here, the structurally identical hexasaccharide to the BOS was found on the *B. petrii* bacterial cells, covalently linked to their LPS molecules. The BOS_LPS_ attached to the inner core oligosaccharide constitutes the main fraction of the clinical *B. petrii* 13363 LPS, whereas environmental *B. petrii* strain 12804 contains LPS with a minor population of the molecules containing the hexasaccharide. This finding of BOS_LPS_ increases the diversity of glycoforms among LPS molecules on the surface of *Bordetella* bacterial cells. No polymeric long O-chains were identified in the *B. petrii* strains. The BOS occupies the position used to be substituted with the terminal trisaccharide, that is characterisitic for *B. pertussis* or O-PS of other *Bordetella* LPS.

Moreover, like other *Bordetella* the BOS was found in the culture media of *B. petrii*, the both clinical isolate and environmental strain. The existence of both structurally identical hexasaccharides in the *B. petrii*, the LPS-bound glycoform BOS_LPS_ and the free glycan BOS suggests that it was partially released from LPS of the *B. petrii* bacterial cells to the bacterial culture.

Some explanation for the association of the hexasaccharide with the *B. petrii* LPS can be provided from gene sequence analyzes of *Bordetella*. With the environmental origin, *B. petrii*, which is also associated with host organisms, perhaps is more closely related to the common ancestor than the pathogenic *Bordetella*. Evolution of the host restricted-pathogens, *B. pertussis, B. bronchiseptica* and *B. parapertussis* was dominated by substantial gene decay. During adaptation to the host, the genes for several metabolic functions have been partially deleted from the genomes of pathogenic *Bordetella* species ^12^. We suppose that *Bordetella* adapting to the host”s immune system changed the surface components through the modification of the terminal elements of the LPS. The presence of the hexasaccharide as the molecule that is produced and released by bacterial cells into surrounding environment in *Bordetella*, but also linked to the core oligosaccharide of the *B. petrii* LPS suggests that the hexasaccharide originates from LPS. The BOS could be a component of the LPS in the early evolution of *Bordetella*, whereas during the adaptation to host as the pathogenic species, the hexasaccharide has been modified and expressed as a secreted molecule. For these reasons, a loss of the hexasaccharide from LPS and presence of the BOS as secreted molecule during the evolutionary process of the bacteria could serve for better adaptation to mammalian hosts. Consequently, the BOS might be one of the factors leading to evolutionary success of pathogenic *Bordetella*.

Other explanation is related to the ability of *B. petrii* to release the hexasaccharide from the bacterial cells during their growth. As a main role of biodiversity of LPS molecules is to avoid the host”s immune defense, the loss of the terminal hexasaccharide from the LPS might be the way to change the epitopes during the host-pathogen interaction. Consequently, another epitopes are exposed, which can impair the immune recognition and in effect prolong the course of the infection. The presence of the glucosamine-1-phosphate in the bacterial culture might confirm the active process of the hexasaccharide release from the *B. petrii* LPS, perhaps during the process of enzymatic phosphorolysis. However, the presence of uridine diphosphate, the activated precursor for the synthesis of cell wall components ^46,47^, and also involved in other metabolic pathways might indicate more complex process in the *B. petrii* growth medium, likely involving the BOS. The detailed mechanisms of the process, synthesis and function of the newly revealed hexasaccharide remain to be investigated.

Except *B. petrii*, the hexasaccharide was found not linked to the LPS molecule but present in the bacterial culture of *Bordetella*. In this form, the BOS might represent one of the factors leading to universal features of this species. Moreover, the amounts of the BOS much higher than in the growth media of *B. petrii* indicate important role of the glycan in typically pathogenic *Bordetella*. It seems that the hexasaccharide released outside the bacterial membrane with relatively high amounts might provide the additional protection from the host immune defense, making the pathogen well-adapted to the mammalian host. The presence of the exo-hexasaccharide in the host-adapted *Bordetella* such as *B. pertussis* suggests importance of the free glycoform in the bacterial pathogenesis and interaction with the host as the clinical strain, required to colonize tissues, adhere to cilia in the respiratory tract, and evade the host immune system. Consequently, it seems that the BOS might play role in adaptive evolution of this genus. We hypothesized that the exo-hexasaccharide turned out to be more functional in the clinical strains, therefore the mechanism of the glycan release from LPS became more effective in the pathogens. The common presence of the hexasaccharide in the *Bordetella* culture suggests its important function similar to this of typical cell-surface exopolysaccharides, as well as the Enterobacterial Common Antigen (ECA) that is shared among all species of *Enterobacteriaceae* ^48^. The polysaccharides play a key role in maintaining the outer membrane permeability barrier as well as by forming biofilm matrix can be involved in bacterial colonization, and protection against environmental and host defenses.

In addition to the potential protective role of the BOS glycoform, the cell-bound BOS_LPS_ might acquire immunogenic properties. The neoglycoconjugate of the *Bordetella* exo-hexasaccharide with tetanus toxoid was able to elicit the antibodies specific for the BOS, in the previous studies ^37^. The anti-BOS antibodies are cross-reactive with *B. petrii* LPS-derived epitopes. Immunogenicity of the BOS-protein carrier conjugate confirms the role of this exo-oligosaccharide as a common element among *Bordetella*. In this aspect, the exo-hexasaccharide shared among all species of *Bordetella* might be considered a potential target for cross-reactive specific antibodies. Whereas, the BOS-containing *B. petrii* LPS could provide antibodies directed against the BOS as well as the inner oligosaccharide core of LPS, making it a potential vaccine antigen against *Bordetella*.

## Methods

### Bacteria and LPS isolation

*B. petrii* strain NCTC 13363 was obtained from the National Collection of Type Cultures, London, UK. *B. petrii* 13363 was the first described clinical strain, from the mandibular osteomyelitis case ^8^. Environmental isolate of *B. petrii* strain DSM 12804 (ATCC BAA-461) was aquired from the Leibniz Institute DSMZ, Germany. The strain was primary isolated from an anaerobic, dechlorinating bioreactor culture enriched from river sediment.

*B. petrii* were grown on charcoal agar containing the defibrinated sheep blood for 48 h at 37°C. Liquid cultures were performed in the Stainer-Scholte medium at 37 °C for 3-5 days with shaking (Infors HT Ecotron, 140 rpm). The growth of bacteria was monitored by biotyping using MALDI-TOF MS Biotyper method. The bacteria were harvested from the medium by centrifugation (8000 × g, 30 min, 4 °C). The LPS were isolated according to the modified hot phenol/water extraction method, by Westphal and Jann ^38^ and purified by ultracentrifugation. LPS samples were resolved by SDS-PAGE and detected by a silver staining ^49^. The supernatant separated from the bacterial cells was used for the exo-hexasaccharide (BOS) isolation.

### Isolation of heteropoly - and oligosaccharides

Isolated *B. petrii* LPS was hydrolyzed with 1.5% acetic acid. Lipid A was separated by centrifugation and the soluble products were further separated by size-exclusion chromatography (HiLoad 16/600 Superdex 30 prep grade). The column was equilibrated with 0.05 M acetic acid.

The eluates were monitored spectrophotometrically and by refractometry. Fractions were collected and checked by NMR and MALDI-TOF mass spectrometry.

### BOS isolation

Supernatant obtained from *Bordetellae* bacterial culture was freeze-dried and subsequently fractionated using size-exclusion chromatography (HiLoad 16/600 Superdex 30 pg, injection 2g/2ml). The eluates were monitored spectrophotometricaly and by refractometry. All fractions were checked by NMR spectroscopy.

### NMR spectroscopy

NMR spectra of the poly- and oligosaccharides in ^2^H_2_O solution were recorded on a Bruker Avance III 600 MHz spectroscope (Bruker). Acetone (δ_H_ 2.225 and δ_C_ 31.05) was used as the internal reference. The signals were assigned by one- and two-dimensional experiments (COSY, TOCSY, NOESY, HMBC, HSQC-DEPT and HSQC-TOCSY). In the TOCSY experiments the mixing times were 30, 60, and 100 ms. The data were acquired and processed using standard Bruker software. The processed spectra were assigned with help of the SPARKY program ^50^. The oligosaccharide ratio was estimated from signal integrals, in the Topspin program.

### Mass spectrometry

MALDI-TOF MS of the poly- and oligosaccharide was performed with the UltrafleXtreme instrument (Bruker Daltonics). 2,5-Dihydroxybenzoic acid was used as the matrix. The OS samples were investigated in the positive and negative ion modes.

### Immunochemical method

Enzyme-linked immunosorbent assay (ELISA) was performed by the method described by Voller *et al* with modification as previously described ^51^. Rabbit anti-exo-hexasaccharide sera were analyzed with LPS (10 μg/ml) as a solid-phase antigen, while goat anti-rabbit IgG conjugated with alkaline phosphatase (Bio-Rad) was used as a second antibody, with *p*-nitrophenyl phosphate as the detection system.

## Supporting information

Koj_Bpetrii_SI

## Data availability

Data generated in this study are provided in the manuscript and its Supplementary Information.

## Supplementary Information

Supplementary Figs. 1-7 and Tables 1-2. Reporting Summary

## Author contributions

SK, KU, TN - concept and design of the work. SK, KU, OR - acquisition, analysis and interpretation of the data. SK - writing the original draft. SK, KU, OR, TN - review and editing the final manuscript. All of the authors approved and contributed to the final version of the manuscript.

