## Supplementary material for "*Bordetella* oligosaccharide (BOS) is associated with lipopolysaccharide of *Bordetella petrii* - the ancestor-related species of the pathogenic *Bordetella*": Koj_Bpetrii_SI: SI_Bpetrii_Fig_Tab_v5_Gal.pdf

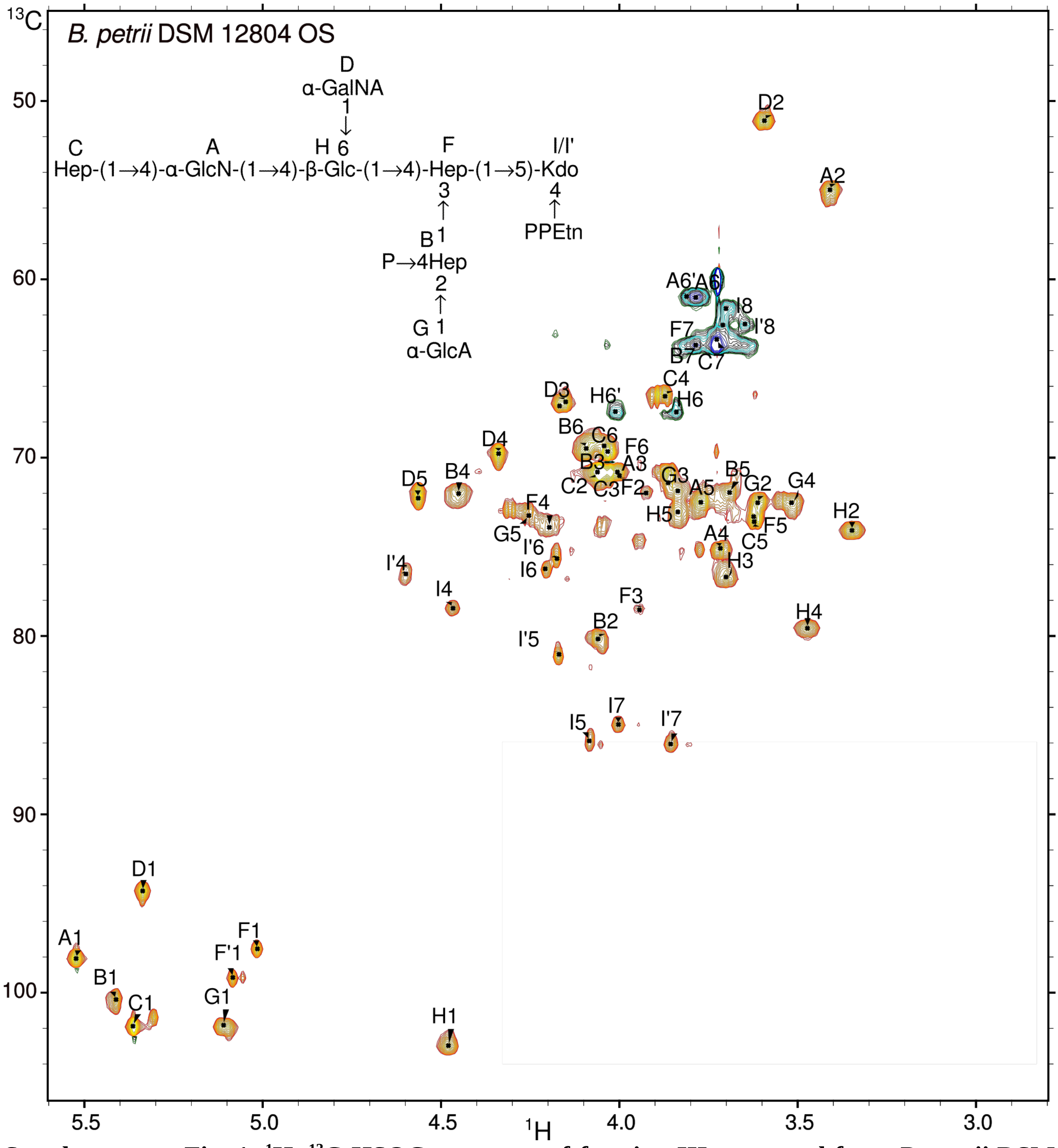


**Supplementary Fig. 1. [1]H, [13]C HSQC spectrum of fraction III separated from *B. petrii* DSM 12804 OS by size-exclusion chromatography.** Letter with a prime sign, e.g. I' is a variant of the residue I that is formed from Kdo after the acid hydrolysis of the oligosaccharide.

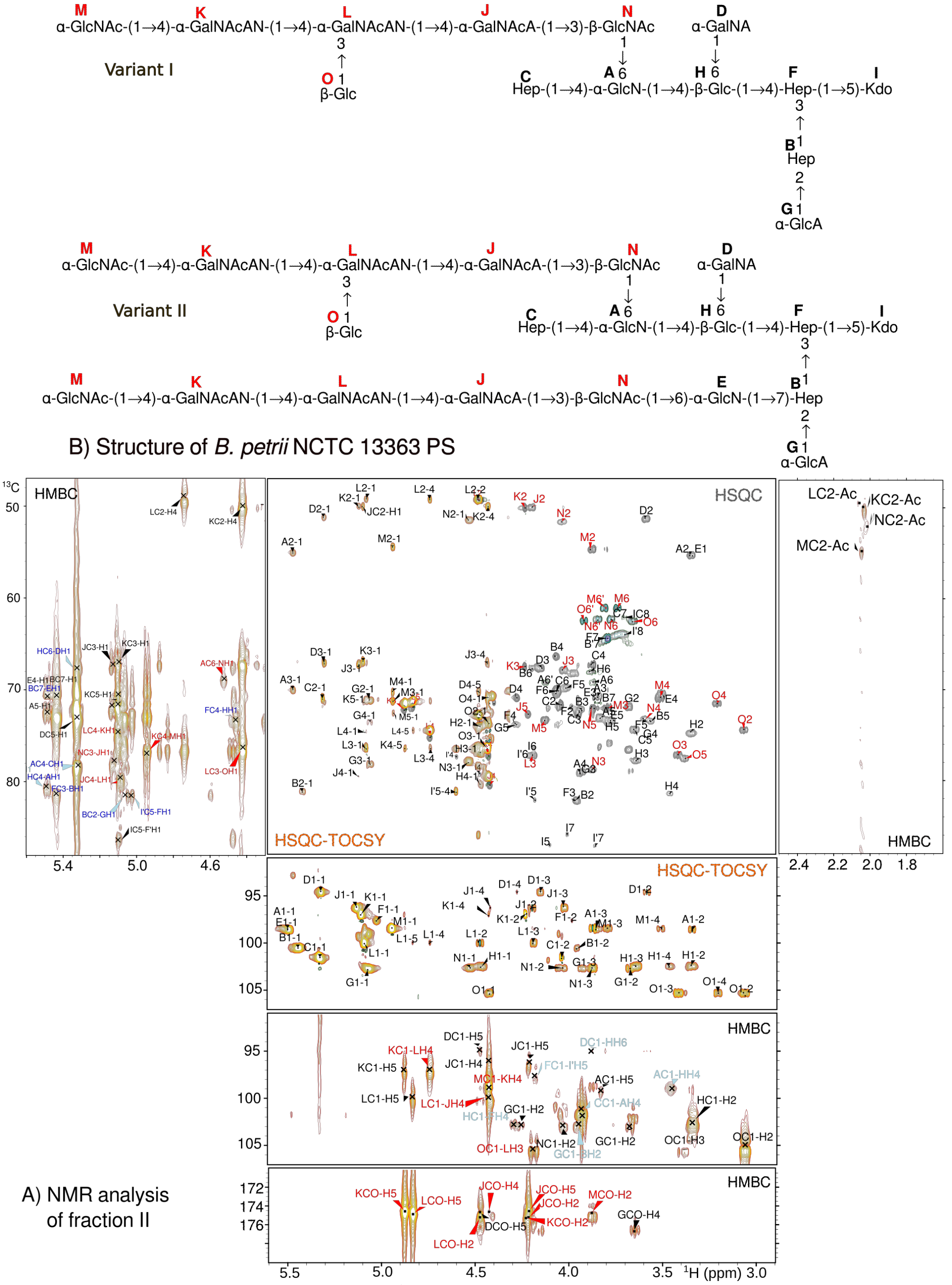


**Supplementary Fig. 2. A) Regions of $^{1}$H,$^{13}$C HSQC-DEPT, HSQC-TOCSY and HMBC spectra of *B. petrii* 13363 fraction II.** Signals indicating the hexasaccharide unit, interglycosidic linkages between the residues of the BOS unit, and between the unit and the core OS are shown in red. B) The structure corresponds to *B. petrii* 13363 PS - core OS with one hexasaccharide unit linked to the residue A. The PS shows heterogeneity regarding the presence of acetyl group on the N residue and the residue E can be absent. As shown on the structures above, two glycoforms are expected, with one unit attached to the A glucosamine and with two units attached to residues A and E, if the E is present.

Two-dimensional $^{1}$H, $^{1}$H COSY, TOCSY, NOESY and $^{1}$H, $^{13}$C HSQC, HMBC, HSQC-TOCSY spectra of the *B. petrii* 13363 fraction II have been analysed (Fig. 3, Fig. S2, Tab. 1). The sequence of the sugar residues was determined in $^1$H, $^1$H NOESY and $^1$H, $^{13}$C HMBC spectra. The following residues of the *B. petrii* 13363 PS were identified:

Residue **A** ($\delta_{H1}/\delta_{C1}$ 5.49/98.6 ppm) was identified as the 4,6-substituted α-Glc*p*N based on large coupling constants between H-2, H-3, H-4, and H-5 in the spin system, as well as the relatively high value of the chemical shift of the C-4 ($\delta_C$ 78.2 ppm) and the C-6 ($\delta_C$ 68.6 ppm) signals. The chemical shift of the C-2 signal ($\delta_C$ 55.1 ppm) indicated a substitution with an amine group. The position 6 of the 4,6-disubstituted α-Glc*p*N residue was identified as a branching point linked to a new element of the *B. petrii* PS.

Residue **B** ($\delta_{H1}/\delta_{C1}$ 5.43/100.5 ppm) was recognized as the 2,7-disubstituted α-L-*glycero*-α-D-*manno*-Hep*p* from the small vicinal couplings between H-1, H-2, and H-3, and the high chemical shifts of the C-2 ($\delta_C$ 81.1 ppm) and C-7 (δC 70.8 ppm) signals. Variant **B'** [→2)-L-α-D-Hep*p*-(1→] is present.

Residue **C** ($\delta_{H1}/\delta_{C1}$ 5.33/101.5 ppm) was assigned as the terminal α-L-*glycero*-α-D-*manno*-Hep*p*.

Residue **D** ($\delta_{H1}/\delta_{C1}$ 5.32/94.6 ppm) was identified as the terminal α-Gal*p*NA based on the large couplings between vicinal protons H-1, H-2, and H-3 and the small couplings among H-3, H-4, and H5, low chemical shift of the C-2 signal ($\delta_C$ 51.1 ppm), the five-proton spin system with high values of the chemical shifts of H-4 ($\delta_H$ 4.27 ppm), H-5 ($\delta_H$ 4.47 ppm), and C-5 ($\delta_C$ 174.6 ppm) resonances characteristic of the aminouronic acid.

Residue **E** ($\delta_{H1}/\delta_{C1}$ 5.50/98.6 ppm) was recognized as the 6-substituted α-Glc*p*N based on low chemical shift of the C-2 signal ($\delta_C$ 55.1 ppm), and the high value of the chemical shift of the C-6 signal ($\delta_C$ 68.6 ppm). This residue could be absent or the position 6 of the →6)-α-Glc*p*N residue is a second branching point of the *B. petrii* PS.

Residue **F** ($\delta_{H1}/\delta_{C1}$ 5.02/97.6 ppm) was characterized as the 3,4-disubstituted α-L-*glycero*-α-D-*manno*-Hep*p* from the high chemical shifts of the C-3 ($\delta_C$ 81.2 ppm) and C-4 ($\delta_C$ 73.0 ppm) signals.

Residues **G** ($\delta_{H1}/\delta_{C1}$ 5.08/102.7 ppm) was recognized as α-Glc*p*A based on the large chemical shifts of H-4 ($\delta_H$ 3.64 ppm), H-5 ($\delta_H$ 4.28 ppm), and C-5 ($\delta_C$ 176.7 ppm) signals.

Residue **H** ($\delta_{H1}/\delta_{C1}$ 4.46/102.5 ppm) was assigned as the 4,6-disubstituted β-Glc*p* based on the large couplings among all protons in the spin system and the high value of the chemical shift of the C-4 ($\delta_C$ 80.5 ppm) and C-6 ($\delta_C$ 67.5 ppm) signals.

Residues **I** and **I'** were assigned as the 5-substituted 4,7-anhydro-3-deoxy-D-*manno*-2-octulofuranose acid forms.

Residue **J** ($\delta_{H1}/\delta_{C1}$ 5.13/96.2 ppm) was identified as the 4-substituted α-Glc*p*NAcA based on high value of the chemical shift of the C-4 signal ($\delta_C$ 79.6 ppm), low chemical shift of the C-2 signal ($\delta_C$ 50.0 ppm), the five-proton spin system with the high values of the chemical shifts of H-4 ($\delta_H$ 4.42 ppm), H-5 ($\delta_H$ 4.22 ppm), and C-5 ($\delta_C$ 174.6 ppm) resonances.

Residue **K** ($\delta_{H1}/\delta_{C1}$ 5.10/96.9 ppm) was identified as the 4-substituted α-Gal*p*NAcAN based on the low chemical shift of the C-2 signal ($\delta_C$ 50.1 ppm), and the high value of the chemical shift of the C-4 signal ($\delta_C$ 76.4 ppm). The presence of amide groups was identified based on proton-nitrogen signal correlation from carbonyl C-5 ($\delta_C$ 174.7 ppm) to H-4 ($\delta_H$ 4.43 ppm) and H-5 ($\delta_H$ 4.88 ppm).

Residue **L** ($\delta_{H1}/\delta_{C1}$ 5.09/99.9 ppm) was identified as the 3,4-substituted α-Gal*p*NAcAN based on low chemical shift of the C-2 signal ($\delta_C$ 49.2 ppm), the high value of the chemical shift of the C-3 ($\delta_C$ 76.4 ppm) and C-4 ($\delta_C$ 74.5 ppm) signals, and the proton-nitrogen signal correlation from carbonyl C-5 ($\delta_C$ 174.9 ppm) to H-4 ($\delta_H$ 4.74 ppm) and H-5 ($\delta_H$ 4.83 ppm).

Residue **M** ($\delta_{H1}/\delta_{C1}$ 4.94/98.4 ppm) was identified as the terminal α-Glc*p*NAc based on low chemical shift of the C-2 signal ($\delta_C$ 54.5 ppm).

Residue **N** ($\delta_{H1}/\delta_{C1}$ 4.53/102.7 ppm) was identified as the 3-substituted β-Glc*p*NAc based on low chemical shift of the C-2 signal ($\delta_C$ 51.5 ppm) and the high value of the chemical shift of the C-3 signal ($\delta_C$ 77.9 ppm). The substitution with acetyl group is partial.

Residue **O** ($\delta_{H1}/\delta_{C1}$ 4.42/105.3 ppm) was identified as the terminal β-Glc*p*.

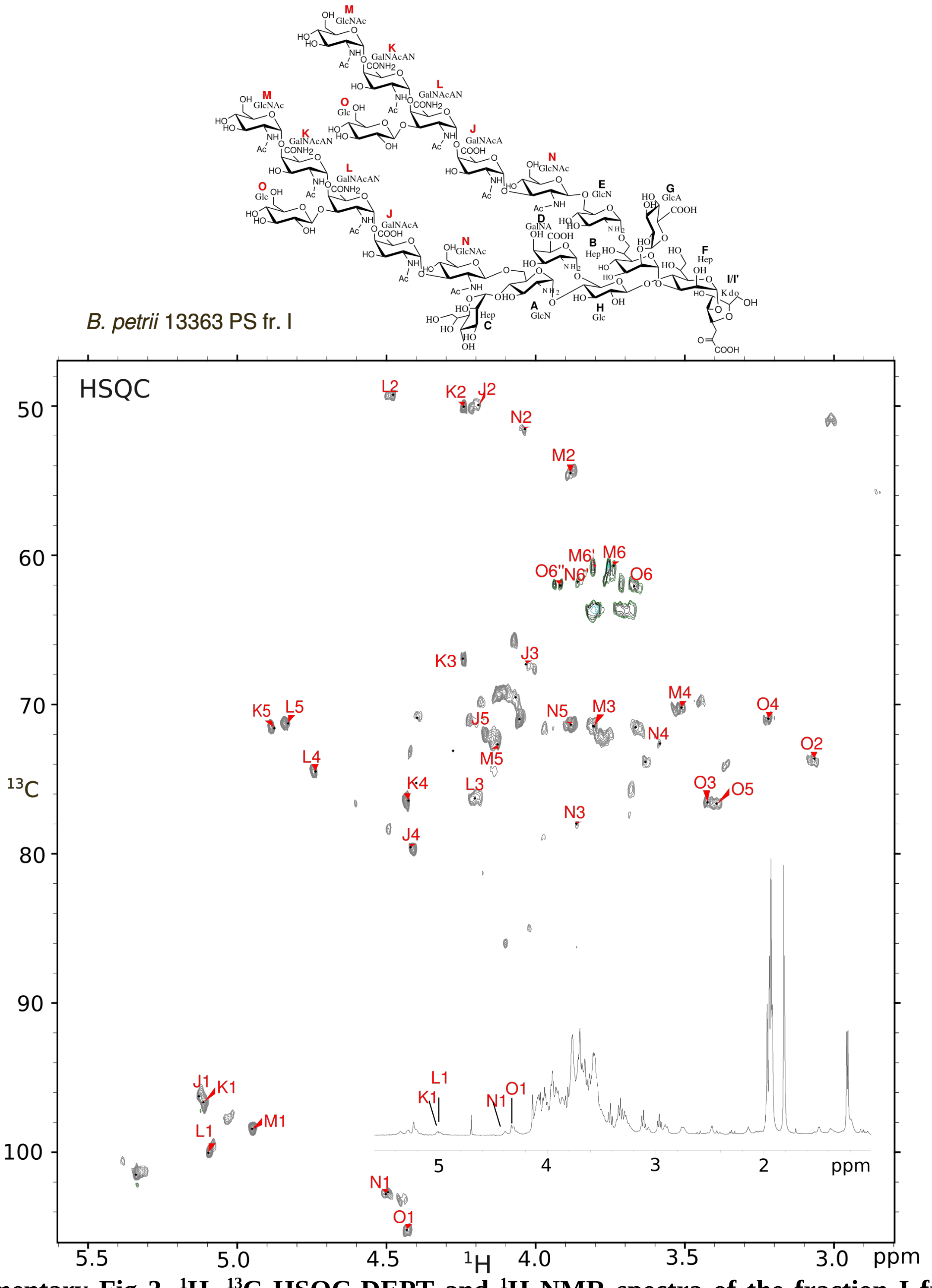


**Supplementary Fig 3. $^{1}H$, $^{13}C$ HSQC-DEPT and $^{1}H$ NMR spectra of the fraction I from *B. petrii* 13363 PS.** The main signals correspond to the residues J-O of the hexasaccharide units. The signals with lower intensities correspond to the core OS (not assigned in the spectra). Two units of the hexasaccharide are likely linked to the core, through A and E residues of the core OS.

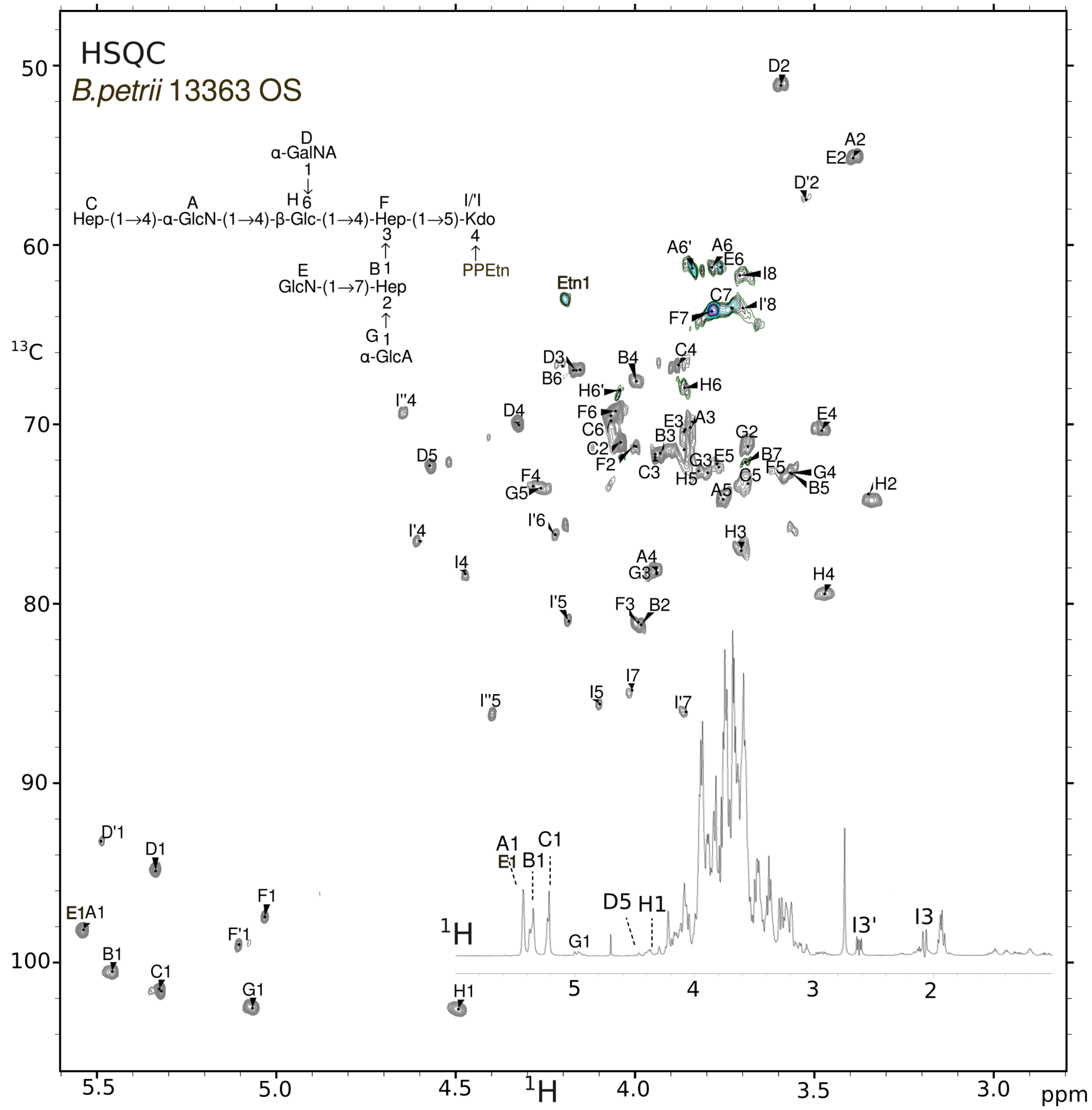


**Supplementary Fig. 4. [1]H, [13]C HSQC-DEPT and [1]H NMR spectra of the fraction IV from *B. petrii* 13363 PS.**

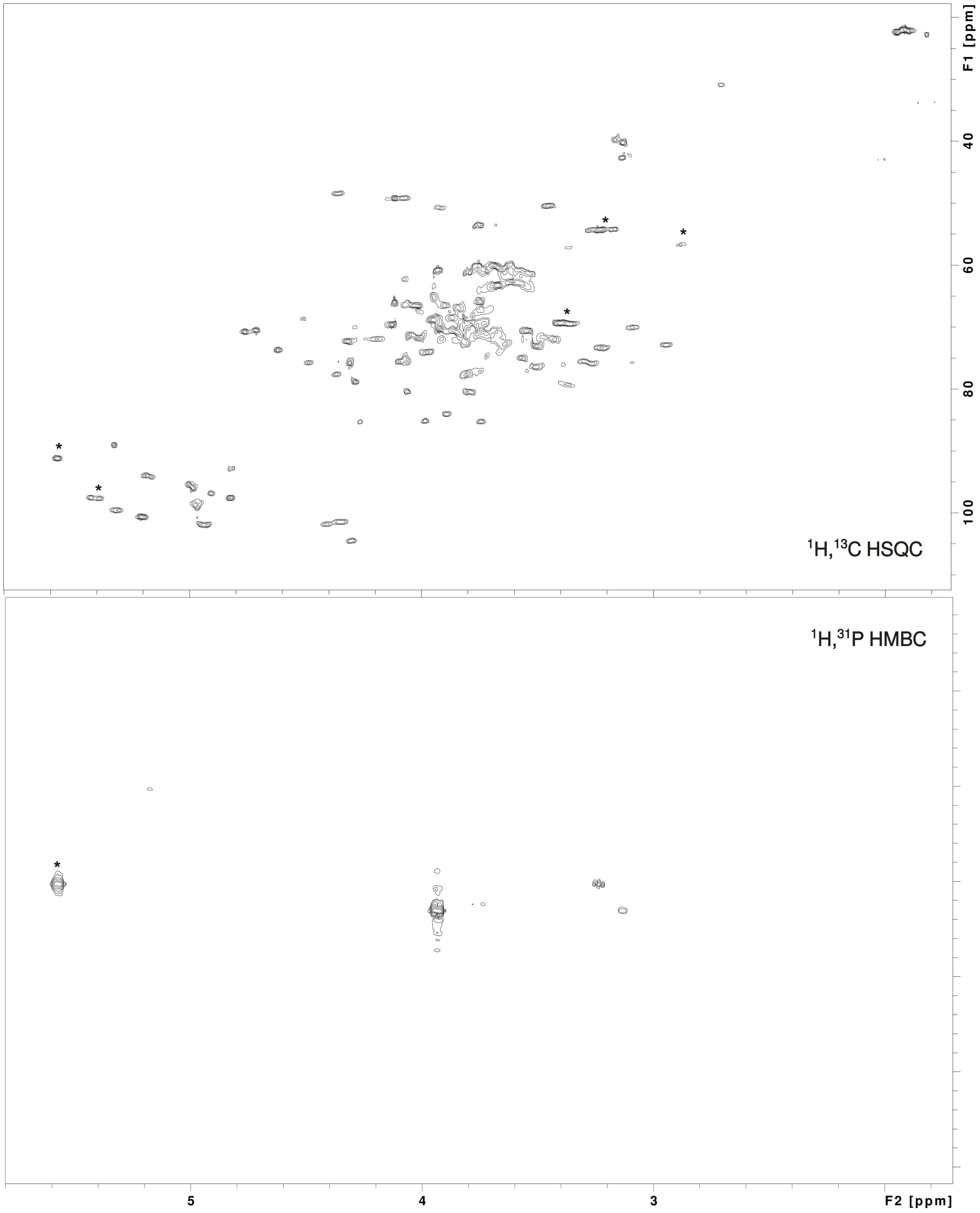


**Supplementary Fig. 5. $^{1}$H, $^{13}$C HSQC-DEPT and $^{1}$H, $^{31}$P HMBC spectra of *B. petrii* 13363 PS.** The signals corresponding to the glucosamines and P-GlcN are indicated by asterics.

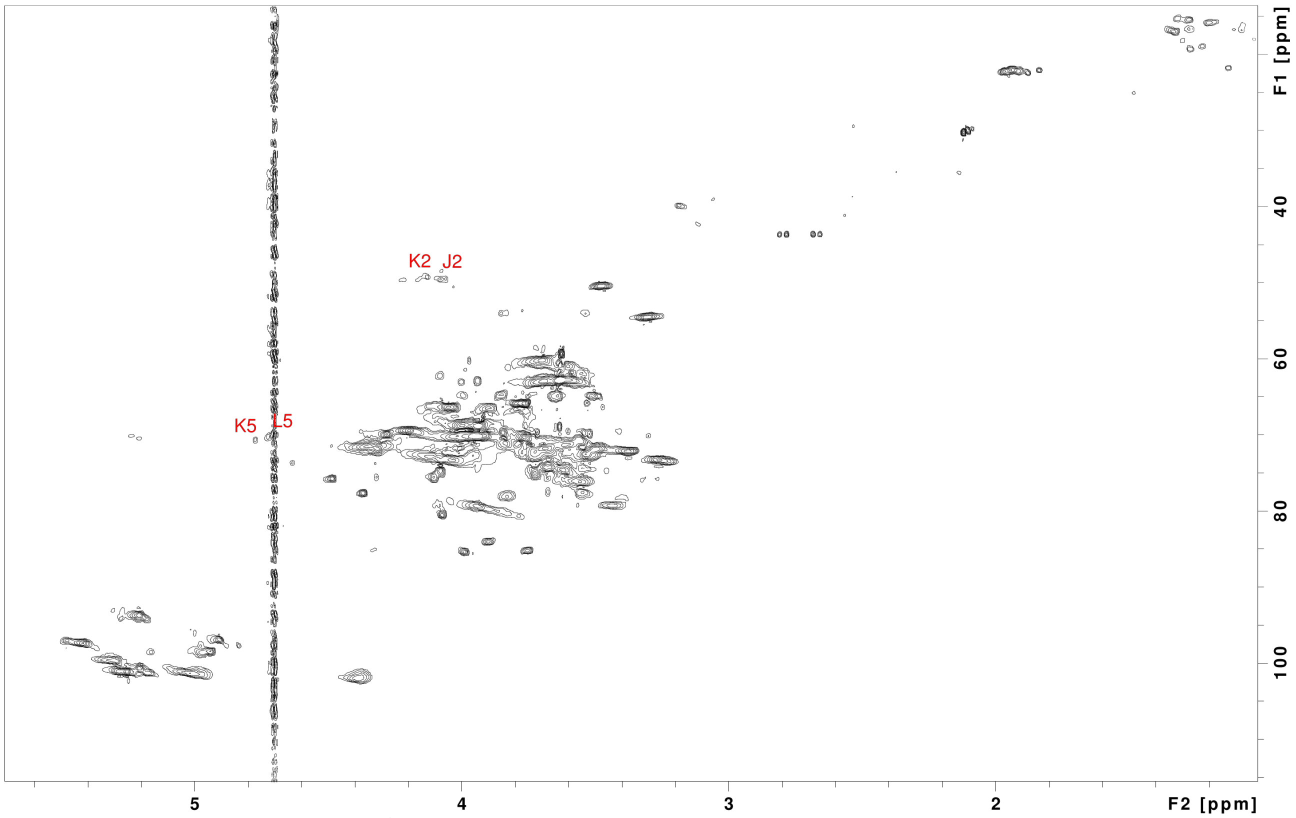

**Supplementary Fig. 6. $^{1}H$, $^{13}C$ HSQC-DEPT spectrum of the fraction II from *B. petrii* 12804 OS.** The signals indicative of the hexasaccharide are assigned in the spectrum.

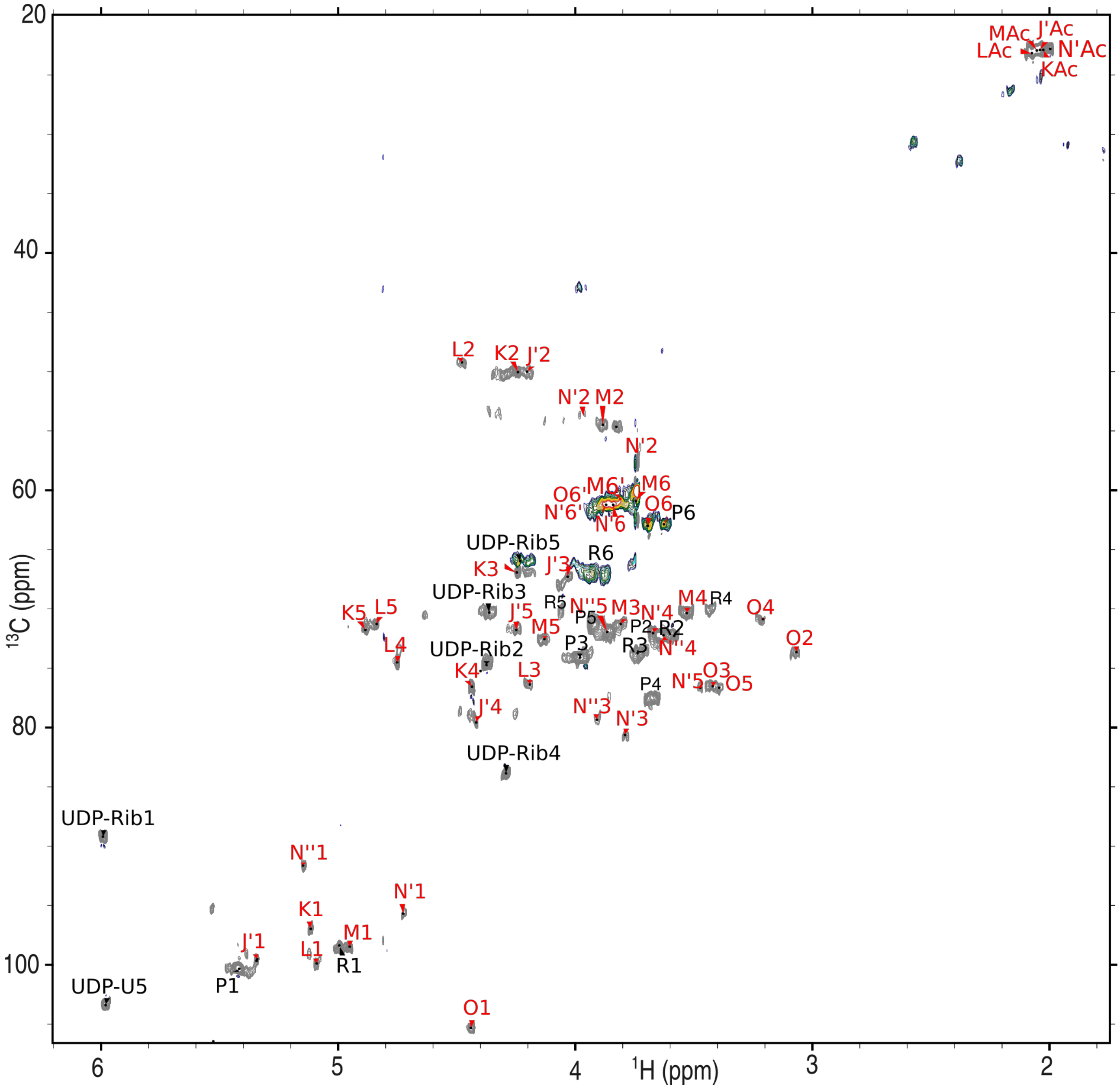


**Supplementary Fig. 7. $^{1}$H, $^{13}$C HSQC spectrum of fraction III from *B. petrii* 13363 bacterial culture.** The spectrum contains signals from the hexasaccharide (red), glucans (P and R) and UDP (black).

**Supplementary Table 1.** $^{1}$**H and** $^{13}$**C chemical shifts of fraction II isolated from *B. petrii* DSM 12804 OS.**

| Sugar residue | Chemical shifts [ppm] | | | | | | | | | |
|---|---|---|---|---|---|---|---|---|---|---|
| | H1<br>C1 | H2<br>C2 | H3<br>C3 | H4<br>C4 | H5<br>C5 | H6, H6'<br>C6 (A) | H7, H7'<br>C7 | H8, H8'<br>C8 (Ac) | Connectivity to<br>Inter-residue atom | $\delta_H$, $\delta_C$ |
| **A** →4,6)-α-D-Glc*p*N-(1→ | 5.52 | 3.41 | 4.00 | 3.72 | 3.77 | 3.80, 3.81 | | | H-C4 | 3.47 |
| | 98.1 | 55.0 | 70.8 | 75.1 | 72.5 | 61.0 | | | H-C4 | 79.5 |
| **B** →2)-L-α-D-Hep*p*4P-(1→ | 5.41 | 4.06 | 4.04 | 4.45 | 3.69 | 4.15 | 3.65, 3.81 | | F-H3 | 3.94 |
| | 100.4 | 80.2 | 70.8 | 72.0 | 72.0 | 69.5 | 63.9 | | F-C3 | 78.5 |
| **C** L-α-D-Hep*p*-(1→ | 5.36 | 4.06 | 3.93 | 3.87 | 3.62 | 4.07 | 3.73 | | A-H4 | 3.72 |
| | 101.9 | 70.8 | 72.0 | 66.6 | 73.6 | 69.8 | 63.4 | | A-C4 | 75.1 |
| **D** α-D-Gal*p*NA-(1→ | 5.34 | 3.59 | 4.15 | 4.34 | 4.56 | | | | H-H6 | 3.84, 4.01 |
| | 94.3 | 51.1 | 66.9 | 69.8 | 72.3 | 173.5 | | | H-C6 | 67.5 |
| **F**→3,4)-L-α-D-Hep*p*-(1→ | 5.02 | 4.00 | 3.94 | 4.20 | 3.62 | 4.03 | 3.67, 3.79 | | | |
| | 97.6 | 71.0 | 78.5 | 73.9 | 73.3 | 69.6 | 63.7 | | I'-C5 | 81.0 |
| **F'**→3,4)-L-α-D-Hep*p*-(1→ | 5.08 | 4.05 | nd | nd | nd | nd | | | | |
| | 99.2 | 71.1 | | | | | | | I-C5 | 85.9 |
| **G** α-D-Glc*p*A-(1→ | 5.11 | 3.61 | 3.86 | 3.52 | 4.26 | | | | B-H2 | 4.06 |
| | 101.8 | 72.5 | 70.9 | 72.5 | 73.2 | 175.3 | | | B-C2 | 80.2 |
| **H** →4,6)-β-D-Glc*p*-(1→ | 4.48 | 3.35 | 3.70 | 3.47 | 3.84 | 3.84, 4.01 | | | F-H4 | 4.20 |
| | 103.0 | 74.1 | 76.7 | 79.6 | 73.0 | 67.5 | | | F-C4 | 73.9 |
| **I**→5)-α-anhydroKdo*f* | | | 2.24 | 4.47 | 4.08 | 4.21 | 4.00 | 3.70 | | |
| | | | 43.4 | 78.4 | 85.9 | 76.3 | 85.0 | 61.7 | | |
| **I'** →5)-α-anhydroKdo*f* | | | 2.23, 2.26 | 4.60 | 4.17 | 4.18 | 3.86 | 3.65<br>3.72 | | |
| | | | 39.7 | 76.5 | 81.0 | 75.7 | 86.0 | 62.5 | | |

**Supplementary Table 2. $^{1}$H and $^{13}$C chemical shifts of fraction IV and fraction I isolated from *B. petrii* NCTC 13363 PS, and exo-hexasaccharide BOS from bacterial culture.**

| Sugar residue | H1<br>C1 | H2<br>C2 | H3<br>C3 | H4<br>C4 | H5<br>C5 | H6, H6'<br>C6 (A) | H7, H7'<br>C7 | H8, H8'<br>C8 (Ac) | Connectivity to<br>Inter-residue atom | $\delta_H$, $\delta_C$ |
|---|---|---|---|---|---|---|---|---|---|---|
| | | | | | **Fr IV** | | | | | |
| **A** →4,6)-α-D-GlcpN-(1→ | 5.54 | 3.39 | 3.85 | 3.94 | 3.75 | 3.79, 3.84 | | | H-C4 | 3.47 |
| | 98.2 | 55.1 | 70.2 | 78.2 | 73.9 | 61.3 | | | H-C4 | 79.5 |
| **B** →2,7)-L-α-D-Hep*p*-(1→ | 5.46 | 3.98 | 3.93 | 4.00 | 3.57 | 4.19 | 3.69, 4.04 | | F-H3 | 3.99 |
| | 100.5 | 81.2 | 71.6 | 67.6 | 72.8 | 67.0 | 72.1 | | F-C3 | 81.1 |
| **C** L-α-D-Hep*p*-(1→ | 5.33 | 4.04 | 3.93 | 3.88 | 3.68 | 4.07 | 3.73 | | A-H4 | 3.94 |
| | 101.5 | 71.1 | 71.6 | 66.7 | 73.3 | 69.8 | 63.4 | | A-C4 | 78.2 |
| **D** α-D-Gal*p*NA-(1→ | 5.34 | 3.59 | 4.15 | 4.32 | 4.57 | | | | H-H6 | 3.86, 4.04 |
| | 94.8 | 51.1 | 67.0 | 70.0 | 72.3 | 173.9 | | | H-C6 | 68.0 |
| **D'** α-D-Gal*p*NA-(1→ | 5.49 | 3.52 | nd | nd | nd | nd | | | | |
| | 93.2 | 57.4 | | | | | | | | |
| **E** α-D-Glc*p*N-(1→ | 5.54 | 3.40 | 3.86 | 3.48 | 3.76 | 3.73, 3.84 | | | | |
| | 98.2 | 55.1 | 70.3 | 70.2 | 72.6 | 61.3 | | | B-C7 | 72.1 |
| **F** →3,4)-L-α-D-Hep*p*-(1→ | 5.03 | 4.00 | 3.99 | 4.29 | 3.60 | 4.06 | 3.79 | | | |
| | 97.5 | 71.2 | 81.1 | 73.4 | 72.9 | 69.5 | 63.7 | | I'-C5 | 81.0 |
| **F'** →3,4)-L-α-D-Hep*p*-(1→ | 5.11 | 4.06 | nd | nd | nd | nd | | | | |
| | 99.1 | 71.1 | | | | | | | I-C5 | 85.6 |
| **G** α-D-Glc*p*A-(1→ | 5.07 | 3.68 | 3.79 | 3.57 | 4.26 | | | | B-H2 | 3.98 |
| | 102.5 | 71.2 | 72.7 | 72.7 | 73.5 | 175.1 | | | B-C2 | 81.2 |
| **H** →4,6)-β-D-Glc*p*-(1→ | 4.49 | 3.35 | 3.70 | 3.47 | 3.82 | 3.86, 4.04 | | | F-H4 | 4.29 |
| | 102.5 | 74.2 | 76.9 | 79.5 | 72.7 | 68.0 | | | F-C4 | 73.4 |
| **I** →5)-α-anhydroKdo*f* | | | 2.83 | 4.47 | 4.10 | 4.22 | 4.01 | 3.71<br>3.78 | | |
| | | | 31.6 | 78.3 | 85.6 | 76.2 | 84.8 | 61.7 | | |
| **I'** →5)-α-anhydroKdo*f* | | | | 4.60 | 4.18 | 4.19 | 3.86 | 3.70<br>3.72 | | |
| | 204.6 | | 39.3 | 76.5 | 81.0 | 75.7 | 86.0 | 63.5 | | |
| I''' Kdo | | | 2.17, 2.73 | 4.65 | 4.40 | | | | | |
| | | 105.3 | 43.7 | 69.3 | 86.2 | | | | | |
| PPEtN | | 4.19 | 3.29 | | | | | | | |
| | | 63.0 | 40.6 | | | | | | | |
| | | | | | **Fr I** | | | | | |
| **J** →4)-α-D-Gal*p*NAcA-(1→ | 5.13 | 4.19 | 4.02 | 4.42 | 4.17 | 174.7 | | 2.04 | N-H3 | 3.86 |
| | 96.2 | 50.0 | 67.3 | 79.6 | 72.0 | | | 22.8, 174.6 | N-C3 | 77.9 |
| **K** →4)-α-D-Gal*p*NAcAN-(1→ | 5.10 | 4.23 | 4.24 | 4.43 | 4.88 | 174.7 | | 2.04 | L-H4 | 4.74 |
| | 96.9 | 50.1 | 66.9 | 76.4 | 71.6 | | | 22.9<br>174.9 | L-C4 | 74.5 |
| **L** →3,4)-α-D-Gal*p*NAcAN-(1→ | 5.09 | 4.48 | 4.19 | 4.74 | 4.83 | 174.7 | | 2.08 | J-H4 | 4.42 |
| | 99.9 | 49.2 | 76.4 | 74.5 | 71.3 | | | 23.1 | J-C4 | 79.6 |

| | | | | | | | | | |
|---|---|---|---|---|---|---|---|---|---|
| | | | | | | | 175.4 | | |
| **M** α-D-Glc*p*NAc-(1→ | 4.94<br>98.4 | 3.88<br>54.5 | 3.80<br>71.3 | 3.51<br>70.2 | 4.13<br>72.7 | 3.74, 3.80<br>60.7 | 2.05<br>22.9<br>174.4 | K-H4<br>K-C4 | 4.43<br>76.4 |
| **N** →3)-β-D-Glc*p*NAc-(1→ | 4.51<br>102.7 | 4.04<br>51.5 | 3.86<br>77.9 | 3.59<br>72.6 | 3.88<br>71.3 | 3.77, 3.85<br>61.9 | 2.03<br>23.1 | A-C6<br>E-C6 | 68.6<br>68.6 |
| **O** β-D-Glc*p*-(1→ | 4.42<br>105.3 | 3.07<br>73.6 | 3.42<br>76.3 | 3.21<br>70.9 | 3.38<br>76.7 | 3.66, 3.92<br>62.1 | | L-H3<br>L-C3 | 4.19<br>76.4 |
| **Fr V** | | | | | | | | | |
| PGlcN | 5.66<br>91.7 | 3.32<br>55.3 | 3.94<br>70.6 | 3.51<br>70.3 | 3.95<br>73.0 | 3.80, 3.84<br>61.1 | | | |
| PEtN | | 4.02<br>61.2 | 3.23<br>41.3 | | | | | | |
| **Exo-hexasaccharide BOS** | | | | | | | | | |
| **J'** →4)-α-D-Gal*p*NAcA-(1→ | 5.34<br>99.5 | 4.20<br>50.0 | 4.03<br>67.3 | 4.42<br>79.6 | 4.25<br>71.8 | 174.2 | 2.04<br>22.8,<br>174.6 | N'-H3<br>N'-C3 | 3.79<br>80.6 |
| **K** →4)-α-D-Gal*p*NAcAN-(1→ | 5.12<br>96.9 | 4.24<br>50.0 | 4.24<br>66.9 | 4.44<br>76.6 | 4.89<br>71.6 | 174.4 | 2.03<br>22.9<br>174.9 | L-H4<br>L-C4 | 4.75<br>74.5 |
| **L** →3,4)-α-D-Gal*p*NAcAN-(1→ | 5.09<br>99.9 | 4.48<br>49.2 | 4.19<br>76.4 | 4.75<br>74.5 | 4.84<br>71.3 | 174.5 | 2.08<br>23.1<br>175.7 | J'-H4<br>J'-C4 | 4.42<br>79.6 |
| **M** α-D-Glc*p*NAc-(1→ | 4.95<br>98.5 | 3.88<br>54.5 | 3.80<br>71.3 | 3.53<br>70.3 | 4.13<br>72.6 | 3.74, 3.81<br>60.8 | 2.05<br>22.9<br>174.4 | K-H4<br>K-C4 | 4.44<br>76.6 |
| **N'** →3)-β-D-Glc*p*NAc | 4.73<br>95.7 | 3.74<br>57.7 | 3.79<br>80.6 | 3.67<br>72.1 | 3.47<br>76.7 | 3.79<br>61.3 | 2.00<br>22.8 | | |
| **N''** →3)-α-D-Glc*p*NAc | 5.15<br>91.6 | 3.97<br>53.6 | 3.91<br>79.4 | 3.63<br>72.3 | 3.87<br>71.9 | 3.66, 3.92<br>61.3 | 2.00<br>22.8 | | |
| **O** β-D-Glc*p*-(1→ | 4.44<br>105.3 | 3.07<br>73.6 | 3.42<br>76.5 | 3.21<br>70.9 | 3.39<br>76.7 | 3.76, 3.94<br>61.5 | | L-H3<br>L-C3 | 4.19<br>76.4 |
| Glucans | | | | | | | | | |
| **P** →4)-α-D-Glc*p*-(1→ | 5.42<br>100.3 | 3.64<br>72.3 | 3.98<br>74.1 | 3.68<br>77.6 | 3.92<br>71.4 | 3.63, 3.69<br>62.9 | | | |
| **R** →6)-α-D-Glc*p*-(1→ | 4.99<br>98.4 | 3.58<br>72.3 | 3.74<br>73.7 | 3.43<br>70.2 | 4.06<br>70.2 | 3.88,3.93<br>67.0 | | | |
| UDP-Rib | 5.99<br>89.2 | 4.37<br>74.7 | 4.36<br>70.3 | 4.29<br>83.9 | 4.23<br>66.0 | | | | |
| UDP-U | | 152.8 | | 166.7 | 5.98<br>103.4 | 7.96<br>142.6 | | | |
